# Influence of iron uptake systems on cefiderocol activity in *Escherichia coli*: at the crossroads of antibiotic resistance and virulence

**DOI:** 10.1101/2025.11.06.687119

**Authors:** Emilie Mallecot, Valentine Berti, Marie Petitjean, Julie Meyer, Signara Gueye, Thibaut Morel-Journel, Olivier Clermont, Laurent Poirel, Erick Denamur, Guilhem Royer, Laurence Armand-Lefevre

**Affiliations:** Assistance Publique-Hôpitaux de Paris (AP-HP), Hôpital Bichat-Claude Bernard, Laboratoire de Bactériologie, Paris, France; Université Sorbonne Paris Nord et Université Paris Cité, INSERM, IAME, Paris, France; University of Fribourg, Faculty of Science and Medicine, Medical and Molecular Microbiology, Fribourg, Switzerland; University of Fribourg, Swiss National Reference Center for Emerging Antibiotic Resistance (NARA), Switzerland; Assistance Publique-Hôpitaux de Paris (AP-HP), Hôpital Henri Mondor, Département de Prévention, Diagnostic et Traitement des Infections, Créteil, France; Université Paris-Est Créteil, IMRB, INSERM U955, Équipe “Virus, Hépatologie, Cancer”, Créteil, France; Assistance Publique-Hôpitaux de Paris (AP-HP), Hôpital Bichat-Claude Bernard, Laboratoire de Génétique Moléculaire, Paris, France

## Abstract

Cefiderocol (FDC) is a new siderophore-conjugated cephalosporin that enters the periplasm *via* iron transport systems. However, the specific contribution of individual iron uptake pathways to FDC activity remains unclear. We investigated the role of 12 iron acquisition systems using several *Escherichia coli* strain collections. FDC MICs were determined in iron-depleted and iron-supplemented media for *E. coli* mutants (Keio and pathogenic island [PAI]-deleted collections) and for clinical wild-type or TEM-producing *E. coli* (WT/TEM-*Ec*) and NDM-producing (NDM-*Ec*) isolates. The distribution of iron uptake genes was assessed in these isolates. In addition, *fec* operon prevalence and genomic location were investigated in *E. coli* genomes from EnteroBase and RefSeq databases. Among Keio and 536-PAI-deleted mutants, only Δ*cirA* and Δ*fiu* (enterobactin system) showed increased FDC MICs (8- and 3-fold, respectively), while Δ*fecA* and Δ*fecB* had lower MICs (3-fold decrease). The *fec* operon, a known extraintestinal virulence factor, was significantly more prevalent in isolates with FDC MICs above median than in those with MICs below (94% *vs*. 38% in WT/TEM-*Ec*; 100% *vs*. 0% in NDM-*Ec*). According to EUCAST breakpoints, 66.7% of *fec*-positive NDM-*Ec* were resistant to FDC while none of the *fec*-negative were. The *fec* operon was found in 46.5% of *E. coli* genomes, including virulent clones (ST131, 77%) and was mostly chromosome-borne (99%). Plasmid-borne *fec* closely resembled that of *Enterobacter hormaechei*, suggesting interspecies transfer. Our findings highlight the role of Fec in reducing FDC susceptibility and promoting resistance in NDM-producing *E. coli*. They challenge the virulence-resistance trade-off demonstrating the ‘liaisons dangereuses’ between iron and antibiotics.

## Introduction

Since the 2000s, the emergence of carbapenemase-producing Enterobacterales (CPE) has become a global public health threat (1). CPE are not only resistant to last-line β-lactams, but also to almost all other classes of antibiotics, leaving few therapeutic alternatives in case of infection. Among these, metallo-β-lactamase (MBL)-producing Enterobacterales that include NDM and VIM enzymes are of particular concern as they represent a major therapeutic challenge due to the lack of effective treatment options. Cefiderocol (FDC) is a novel class of β-lactam, corresponding to a siderophore-conjugated cephalosporin molecule active on difficult-to-treat Gram negative bacilli including CPEs (2). The chemical structure of FDC is close to cefepime and ceftazidime and contains a catechol-type siderophore. Its activity is based on a “Trojan horse” strategy that uses active catechol systems by mimicking siderophore molecules to facilitate the entry of FDC into the periplasmic space. Once inside, FDC binds to penicillin-binding proteins (PBP) and then inhibits peptidoglycan synthesis. Therefore, FDC has become a valuable option in the treatment of infections caused by CPE including those that produce MBL. Under physiological conditions, free iron concentrations in human tissues are extremely low (∼10^-24^ M as ferric ion Fe^3+^) (3). Iron is mainly present as an insoluble form and is strongly bound to proteins such as transferrin to limit its toxicity and bacterial proliferation. It is therefore essential for bacteria to synthesize siderophores and iron uptake systems to obtain the iron needed to survive and multiply in iron-deficient conditions (4). Indeed, iron uptake systems are considered to be among the most important virulence factors in bacteria. *Escherichia coli* have numerous iron uptake systems enabling transport of Fe^3+^, Fe^2+^ and heme. Fe^3+^ transport systems include siderophores such as enterobactin (Ent), salmochelin (Iro), yersiniabactin (Ybt) or aerobactin (Iuc) associated with receptors FepA, Fiu, CirA, IutA, IroN and FyuA, high affinity iron transporters such as the ferric di-citrate transporter FecA and the ferrichrome transporter FhuA/FhuE. Fe^2+^ is incorporated through non-specific porins (OmpC/OmpF) and transported by the Feo, Efe and Sit systems. Finally, heme uptake systems involve ChuA and Hma. (5,6).

Since its introduction, several mechanisms associated with decreased susceptibility or resistance to FDC have been reported. Certain β-lactamases, particularly MBLs such as NDM, have been linked to reduced FDC activity, with increased MICs reaching up to 256 mg/L (7,8). Mutations in genes encoding iron acquisition systems, notably *cirA* and *fiu* have also been described and are associated with increased FDC MICs (9,10). However, the influence of siderophores and other iron capture systems in *E. coli* has not been thoroughly investigated. In this study, we aimed to elucidate the role of various iron uptake systems in FDC activity using i) well-characterized collections of *E. coli* mutant strains lacking β-lactamase production, and ii) a collection of clinical isolates producing multiple acquired antibiotic resistance mechanisms, including MBL-producers.

## Materials and Methods

Several collections of strains were used in this study

### Keio knockout collection

The Keio collection is a library of single-gene deletion mutants derived from the *E. coli* K-12 strain BW25113 exhibiting a wild-type antibiotic susceptibility pattern (11). For this study, seventeen mutants with deletions in genes involved in iron uptake (Δ*cirA*, Δ*fiu*, Δf*epA*, Δ*fepB*, Δ*fepC*, Δ*entB*, Δ*fhuE*, Δ*fhuB*, Δ*fhuD*, Δ*fecA*, Δ*fecB*, Δ*fecC*, Δ*ompF*, Δ*efeU*, Δ*efeO*, Δ*feoA*, Δ*feoB*) were selected. The wild-type K-12 BW25113 strain (K-12-WT) was used as control. The presence of the kanamycin resistance cassette in Keio mutants was confirmed by PCR using specific primers (Thermo Fisher Scientific, Waltham, MA, USA), followed by gene-specific PCRs to verify each deletion; control PCRs were also performed on the K-12-WT strain.

### 536-PAI-deleted mutants

A collection of mutants derived from the uropathogenic *E. coli* 536 strain (Hacker *et al*. 1983) was previously constructed by single or multiple deletions of pathogenicity-associated islands (PAIs) (12). For this study, three mutants with deletion of PAI that include genes involved in iron uptake systems were selected: ΔPAI-III (salmochelin and heme receptor HmuR), ΔPAI-IV (yersiniabactin) and PAI-dTTP (all PAIs deleted). The wild-type *E. coli* 536 strain (536-WT) was used as the control.

### *E. coli* clinical isolate collections

Clinical isolates included 65 non duplicated wildtype or TEM-1 (narrow-spectrum β-lactamase) producing *E. coli* (WT/TEM-*Ec*) previously characterized by Royer *et al*. (13) and 24 non redundant and non-clonal NDM-1-producing *E. coli* (NDM-*Ec*) isolated from patients hospitalized at the Bichat-Claude Bernard University Hospital. NDM-*Ec* were mainly from rectal or throat screening samples (n=20), the remaining were from urinary (n=2), respiratory (n=1) and blood (n=1) samples. These NDM-*EC* isolates were selected according to their plasmid harboring the *bla*_NDM-1_, which were either IncFK-pNDM_Epi1 (n=16) or IncC-pCMY-4 (n=9), already described (14) or available in Supplementary Materials. Illumina whole genome sequences of all clinical isolates were available.

### Cefiderocol susceptibility testing

Minimum inhibitory concentrations (MICs) of FDC were determined using the UMIC^®^ Cefiderocol kit (Biocentric, Brandol, France) in iron-depleted cation-adjusted Mueller Hinton Broth (ID-CAMHB, Biocentric), according to EUCAST 2023 guidelines. Each MIC was determined at least twice. Additional measurements were determined if the results differed by more than one dilution.

### Cefiderocol susceptibility testing in iron-supplemented MH broth

To assess the impact of iron concentration on FDC MIC, 8 strains from the Keio (Δ*cirA*, Δ*fiu*, Δ*entB*, Δ*fecA*, Δ*ompF*) and PAI (ΔPAI-III, ΔPAI-IV and PAI-dTTP) collections were selected as well as both wildtype strains (K-12-WT and 536-WT). ID-CAMHB were enriched with ammonium iron (III) citrate (F-5879-100G, Sigma-Aldrich, France) to reach 9 iron concentrations: 0.01, 0.05, 0.1, 0.5, 1, 5, 10, 50, 100 mg/L. FDC MIC of each strain was determined in duplicate at each of the 9 iron concentrations.

### Search for iron uptake systems on genomes

When analyzing genomes of clinical isolates, resistance and virulence genes were searched using Abricate v1.0.0 using ResFinder4 (coverage and identification cut-off fixed at 90% and 70% respectively) and a custom database respectively. The latter included genes or gene operons related to iron uptake system, namely *chu, hmA, ent, cirA, fiu, fep, fes, ire, iro, irp, ybt, fyuA, iuc, iut, fhu, fec, ompC, ompF, efe, feo, sit, tonB, exb, baeS, envZ*. Sequences were found in the NCBI (https://www.ncbi.nlm.nih.gov) and UniProt (https://www.uniprot.org) database.

Additionally, the presence of the *fec* operon was screened in the genomes of the 70,301 *E. coli* strains indexed in Enterobase (https://enterobase.warwick.ac.uk/) (15) and the complete genomes of 2,302 *E. coli* strains available in RefSeq (https://www.ncbi.nlm.nih.gov/refseq) (13). The location of the *fec* operon (plasmidic or chromosomal) was determined for the genomes of *E. coli* from RefSeq and of three clinical NDM-*Ec* isolates, all generated using long-read approach. Plasmid typing was performed using Abricate with the Plasmidfinder database (16) applying identity and coverage thresholds of 80%. Finally, the genetic environment of the *fec* operon in all plasmid sequences carrying a complete operon (*fecABCDE*) were analyzed and compared with Clinker (17) using a 20-kbp region downstream and upstream of *fecA*, including the plasmid previously described by Polani *et al*. (18) (accession number JN233704.1).

### Statistical analysis

R (https://www.R-project.org/) and RStudio software (Posit Software, PBC) were used to perform biostatistics analysis and to create graphic representations. FDC CMIs of each Keio and 536-PAI-deleted mutant were compared to corresponding wild type strains, K-12-WT and 536-WT respectively, using Wilcoxon test. The effect of iron on the FDC MICs of the selected mutants could be analyzed and represented by the LOESS (Locally Estimated Scatterplot Smoothing) regression method. Student and Fisher tests were carried out to determine the significant associations between the iron transport genes and the MICs of the clinical strains. P-values<0.05 were considered as statistically significant.

For each of the WT/TEM-*Ec* and NDM-*Ec* collections, a principal component analysis (PCA) was performed on a matrix of presence in each isolate of operons related to the iron uptake systems listed above. An operon was considered present (noted 1 in the matrix) if more than half of its genes were detected, and absent (noted 0) otherwise. For each analysis, the set of principal components explaining more than 90% of the observed variance were used as the explanatory factors of a linear regression on the Log_2_ of the observed MICs. These two regressions provided estimates of the change in MIC along the principal components for the isolates of the two collections.

## Results

### Cefiderocol activity on Keio and 536-PAI-deleted mutants

The median FDC MIC of the K-12-WT was 0.09 +/-0.04 mg/L. Among the 17 single-deleted mutants from the Keio collection, only four showed an MIC to FDC that was significantly different from that of the K-12-WT strain. The MIC of Δ*cirA and* Δ*fiu* mutants were 0.75 mg/L and 0.25 mg/L, respectively representing 8-fold and 3-fold increases compared to the K-12-WT (p=0.025 and 0.027). Conversely, the MICs of the Δ*fecA* and Δ*fecB* mutants were both 0.03 mg/L, corresponding to a 3-fold decrease relative to K-12-WT (p=0.019 and 0.047). Although not statistically significant, the MIC of the Δ*ompF* (0.19 mg/L) tended to be higher than that of the K-12-WT strain (p=0.086) (Figure 1A).

**Figure 1.**
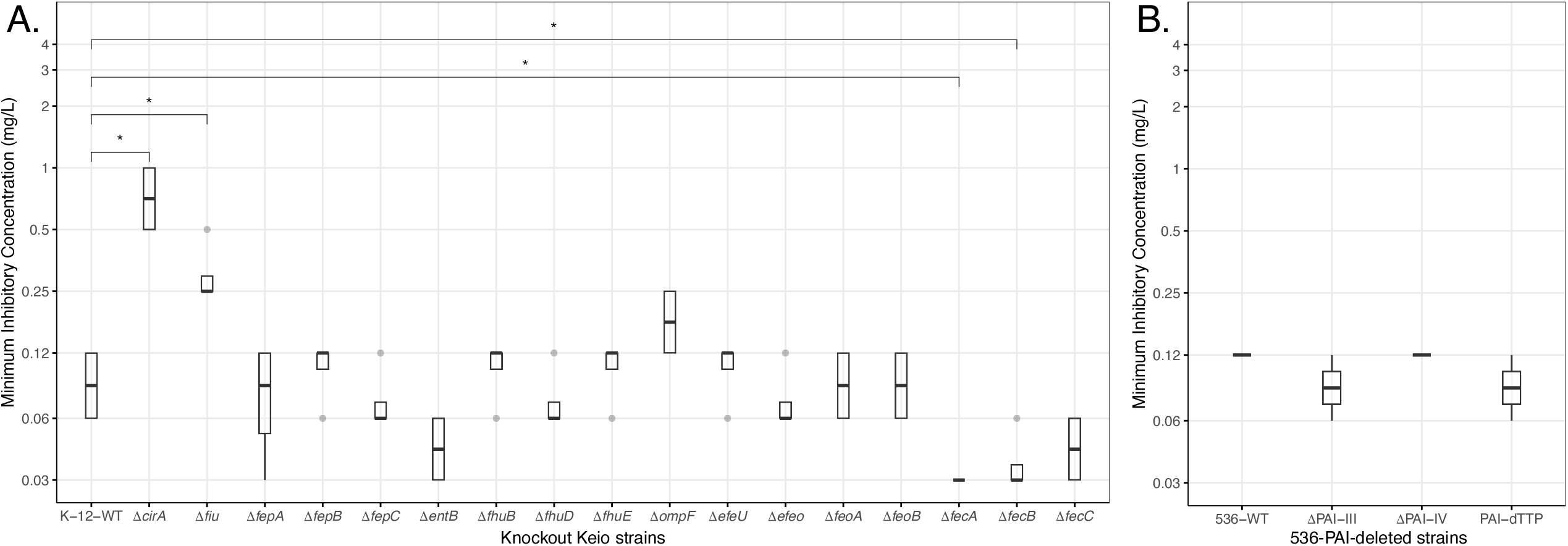
Cefiderocol (FDC) minimum inhibitory concentrations (MIC) of Keio and 536-PAI-deleted mutants in iron-depleted media (ID-CAMHB) for (A) 17 Keio mutants (Δ*cirA*, Δ*fiu*, Δ*fepA*, Δ*fepB*, Δ*fepC*, Δ*entB*, Δ*fhuB*, Δ*fhuD*, Δ*fhuE*, Δ*ompF*, Δ*efeU*, Δ*efeO*, Δ*feoA*, Δ*feoB*, Δ*fecA*, Δ*fecB*, Δ*fecC*) and (B) 3 PAI-deleted mutants (ΔPAI-III, ΔPAI-IV, PAI-dTTP). MICs median for each mutant was compared to those of the K-12-WT (A) or 536-WT (B) strains. * indicate p<0.05

Median FDC MICs of the ΔPAI-III (salmochelin and heme receptor HmuR), ΔPAI-IV (yersiniabactin) and PAI-dTTP (all 536-PAI-deleted) mutants ranged from 0.06 to 0.125 mg/L with no difference compared to the 536-WT strain (Figure 1B).

### Cefiderocol activity on Keio and 536-PAI-deleted mutants with increasing iron concentration

The influence of the different iron uptake systems under increasing iron concentrations were measured for selected mutant strains using ID-CAMHB supplemented with iron (0 to 100 mg/L). The Δ*cirA*, Δ*fiu* and Δ*fecA* mutants were included based on their significant MIC changes observed in iron-depleted media. All 536-PAI-deleted mutants were tested to evaluate the contribution of the salmochelin and yersiniabactin systems. The Δ*entB* mutant, a catecholate-type siderophore naturally produced by *E. coli*, was also included. Finally, the Δ*ompF* mutant was tested due to its slightly decreased FDC susceptibility in iron-depleted conditions, despite not being directly involved in siderophore-mediated iron uptake but rather enabling the diffusion of FDC. The FDC MICs of all tested strains significantly increased with rising iron concentrations. (p<0.05) (Figure 2A, 2B and Table S1). For the K-12-WT, Δ*ompF and* Δ*entB* strains, MICs of FDC increased 4-, 4- and 6-fold, respectively, reaching respective final values of 0.5, 0.38 and 0.31 mg/L under 100 mg/L iron concentrations (regression slopes of 1.6×10^-3^, 3.0 ×10^-3^ and 2.4×10^-3^ respectively). The Δ*cirA* mutant showed the most marked increase in FDC MIC, rising 12-fold to reach 3 mg/L at 50 mg/L iron, with a regression slope of 2.6 × 10^−2^. Similarly, the FDC MIC of the Δ*fiu* mutant increased 3-fold, to 0.5 mg/L (regression slope = 3.3×10^-3^). In contrast, the Δ*fecA* mutant showed a limited increase (regression slope = 4.7 ×10^-4^), with MICs remaining stable at 0.03 mg/L up to 5 mg/L and slightly increasing to 0.09 mg/L at 100mg/L (Figure 2A and Table S1).

**Figure 2.**
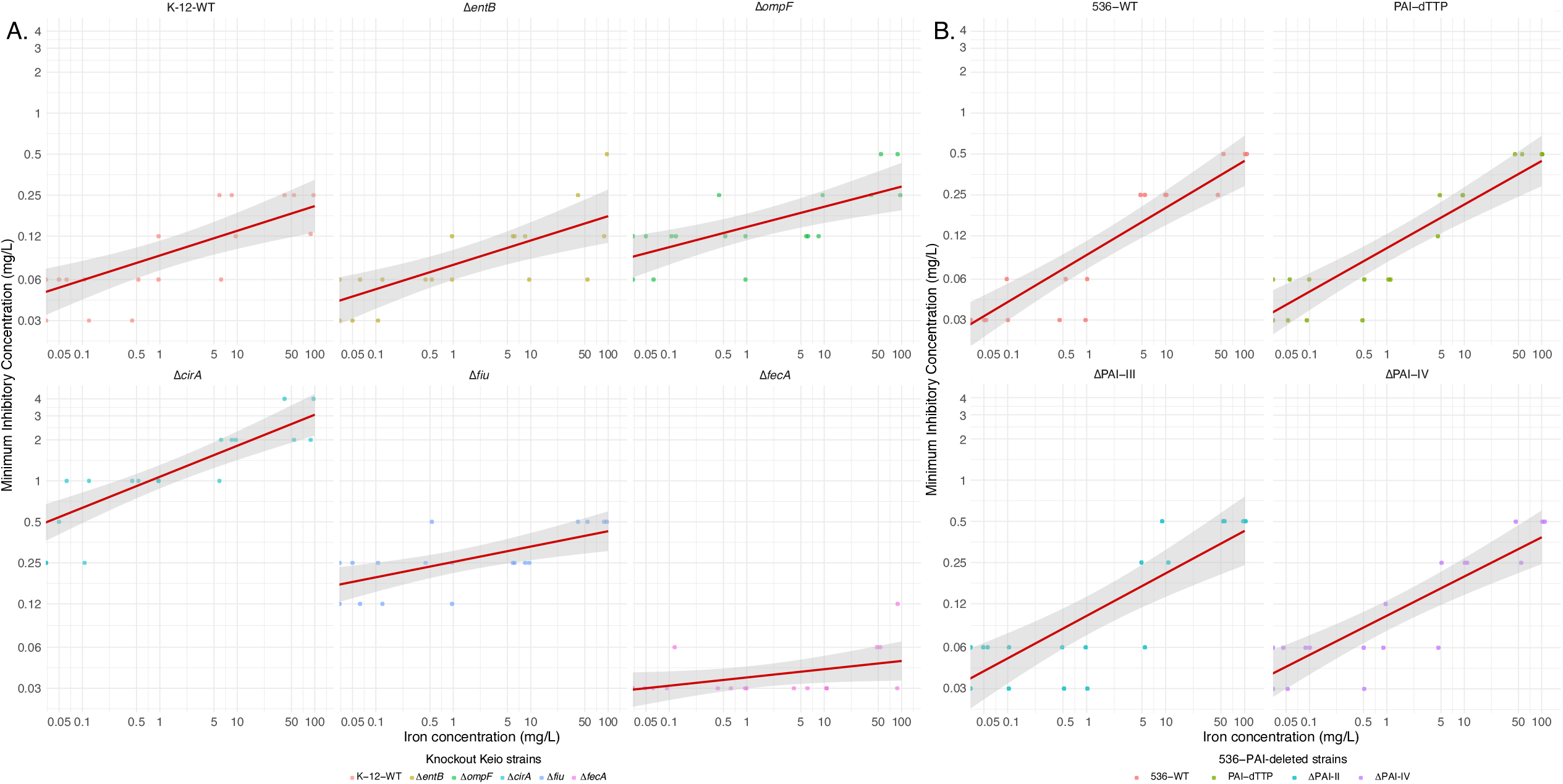
Locally Estimated Scatterplot Smoothing (LOESS) representation of Cefiderocol (FDC) minimum inhibitory concentrations (MIC) of Keio (A) and 536-PAI-deleted (B) mutants with increasing iron concentration. FDC MICs were measured in ID-CAMHB media supplemented with iron from 0 to 100 mg/L. MICs were determined for (A) five Keio mutants (Δ*cirA*, Δ*fiu*, Δ*entB*, Δ*ompF*, Δ*fecA*) and (B) three 536-PAI-deleted mutants (ΔPAI-III, ΔPAI-IV, PAI-dTTP) at each iron concentration. K-12-WT (A) or 536-WT (B) strains were tested as control.

For 536-PAI-deleted mutants, no significant differences in MIC were observed compared to 536-WT strain, trends with regression slopes ranging from 4.6×10^-3^ to 5.0×10^-3^ *vs* 4.7×10^-3^ (Figure 2B and Table S1).

### MICs of FDC on WT/TEM-*Ec* and NDM-*Ec* clinical isolates and prevalence of iron uptake systems

The FDC MICs of the 65 WT/TEM-*Ec* isolates ranged from 0.03 to 4 mg/L with a median of 0.25 mg/L. The FDC MICs of the 24 NDM-*Ec*, MICs ranged from 0.5 to 12 mg/L with a median of 1.75 mg/L. No significant difference in FDC MICs was observed based on the NDM-encoding plasmid (Figure 3). According to EUCAST breakpoint (2 mg/L), 98% (64/65) of the WT/TEM-*Ec* strains and 67% (16/24) of the NDM-*Ec* were categorized as susceptible to FDC.

**Figure 3.**
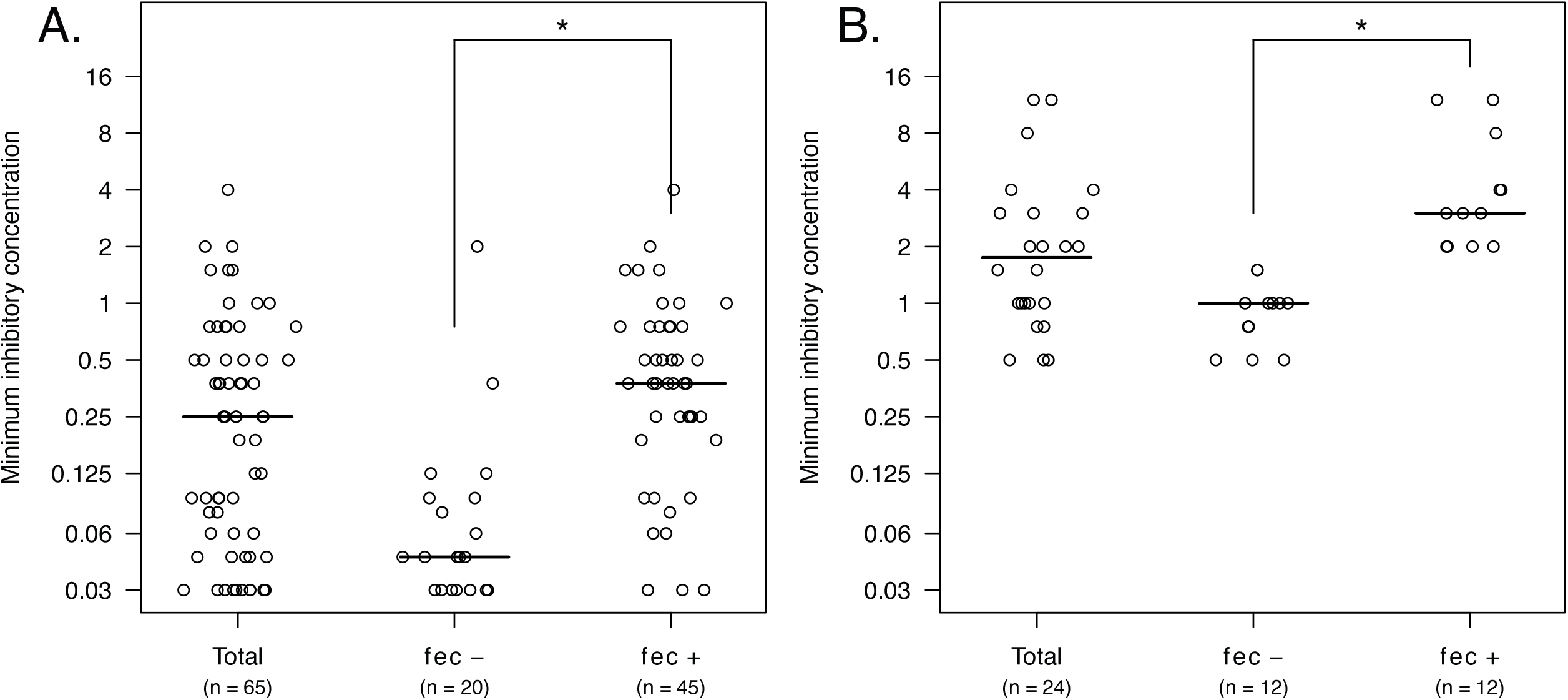
Distribution of FDC MICs among all, *fec-*positive and *fec-*negative isolates from the two collections of *E. coli* clinical isolates: wild-type or TEM-producing *E. coli* (WT/TEM-*Ec)* (A) and NDM-producing *E. coli* (NDM-*Ec*) (B); FDC breakpoint threshold from EUCAST (= 2 mg/L) indicated with a dotted line.

The prevalence of the 12 iron uptake systems in WT/TEM-*Ec* and NDM-*Ec* isolates is presented in Table 1. For each group, their prevalence was compared between isolates with FDC MICs < median or ≥ median. The only iron uptake system significantly associated with higher MICs in both WT/TEM-*Ec* and NDM-*Ec* was the *fec* operon. Its prevalence differed from 94% (34/36) to 38% (11/28) in WT/TEM-*Ec* isolates with MICs above and below the median, respectively, and from 100% (12/12) to 0% (0/12) in NDM-*Ec*. The median MIC of *fec*-positive vs *fec*-negative isolates were 0.38 and 0.05 mg/L (p=0.005) for WT/TEM-*Ec* respectively and 3 and 1 mg/L for NDM-*Ec* (p=0.005) respectively (Figure 3). According to the EUCAST breakpoint, 66.7% (8/12) of *fec*-positive NDM-*Ec*, were resistant to FDC, while none of the *fec*-negative were (p=0.001). The aerobactin system (*iuc/iut*) was also more prevalent in WT/TEM-*Ec* with MIC above the median (64%, 23/36) than below (28%, 8/29). However, this appeared to be linked to co-occurrence with *fec* as 95.7% (22/23) of *iuc-*positive isolates also carried the *fec* operon. Moreover, the MICs of *iuc*-positive/*fec*-positive vs *iuc*-negative/*fec*-positive WT/TEM-*Ec* did not differ significantly (p=0.984).

**Table 1.** Prevalence of iron uptake systems in the two collections of *E. coli* clinical isolates (WT/TEM-*Ec* and NDM-*Ec*). Isolates of each collection were divided into two groups according to their FDC MIC relative to the median MIC of the corresponding collection.

| genes/gene operon | Cefiderocol MICs (mg/L) |  |  |  |  |  |  |  |
| --- | --- | --- | --- | --- | --- | --- | --- | --- |
|  | WT/TEM-*Ec* |  |  |  | NDM-*Ec* |  |  |  |
|  | Total<br>n (%) | <0,25 mg/L<br>n (%) | ≥0,25 mg/L<br>n (%) | P-value | Total<br>n (%) | <1.75 mg/L<br>n (%) | ≥1.75 mg/L<br>n (%) | P-value |
| **Strains total n (%)** | 65 | 29/65 (46) | 36/65 (54) |  | 24 | 12/24 (50) | 12/24 (50) |  |
| **<u>Heme transport systems</u>** | - |  |  |  |  |  |  |  |
| *chu* | 28/65 (43) | 18/29 (62) | 20/36 (56) | 0.62 | 8/24 (33) | 6/12 (50) | 4/12 (33) | 0.68 |
| *hma* | 12/65 (19) | 6/29 (21) | 6/36 (17) | 0.75 | 6/24 (25) | 4/12 (33) | 2/12 (17) | 0.64 |
| **<u>Ferric transport systems</u>** | - |  |  |  |  |  |  |  |
| **-TonB complex** (tonB/exbB/exbD) | 65/65 (100) | 29/29 (100) | 36/36 (100) | 1 | 24/24 (100) | 12/12 (100) | 12/12 (100) | 1 |
| **-Siderophore systems** |  |  |  |  |  |  |  |  |
| enterobactin (*ent/fep/fiu/cirA*) | 65/65 (100) | 29/29 (100) | 36/36 (100) | 1 | 24/24 (100) | 12/12 (100) | 12/12 (100) | 1 |
| salmochelin (*iro*) | 28/65 (43) | 12/29 (41) | 16/36 (44) | 1 | 12/24 (50) | 5/12 (42) | 7/12 (58) | 0.64 |
| aerobactin (*iuc/iutA*) | 31/65 (48) | 8/29 (28) | 23/36 (64) | <0.05 | 8/24 (33) | 2/12 (17) | 6/12 (50) | 0.19 |
| yersiniabactin (*irp/ybt/fyuA*) | 34/65 (52) | 14/29 (48) | 20/36 (56) | 0.62 | 12/24 (50) | 5/12 (42) | 7/12 (58) | 0.64 |
| putative siderophore system (*ireA*) | 13/65 (20) | 5/29 (17) | 8/36 (22) | 0.76 | 2/24 (8) | 0/12 (0) | 2/12 (17) | 0.48 |
| **-Iron transporters** |  |  |  |  |  |  |  |  |
| citrated-base hydroxamate (*fhu*) | 47/65 (72) | 21/29 (72) | 26/36 (72) | 1 | 24/24 (100) | 12/12 (100) | 9/12 (100) | 0.23 |
| di-citrate (*fec*) | 45/65 (69) | 11/29 (38) | 34/36 (94) | <0.05 | 12/24 (50) | 0/12 (0) | 12/12 (100) | <0.05 |
| **<u>Ferrous transporter systems</u>** |  |  |  |  |  |  |  |  |
| *efe* | 35/65 (54) | 16/29 (55) | 19/36 (53) | 1 | 24/24 (100) | 12/12 (100) | 12/12 (100) | 1 |
| *feo* | 65/65 (100) | 29/29 (100) | 36/36 (100) | 1 | 24/24 (100) | 12/12 (100) | 12/12 (100) | 1 |
| *sit* | 36/65 (55) | 17/29 (59) | 19/36 (52) | 0.80 | 17/24 (71) | 9/12 (75) | 8/12 (67) | 1 |
| **<u>Non-specific porins</u>** |  |  |  |  |  |  |  |  |
| *ompF* | 65/65 (100) | 29/29 (100) | 36/36 (100) | 1 | 24/24 (100) | 12/12 (100) | 12/12 (100) | 1 |
| *ompC* | 65/65 (100) | 29/29 (100) | 36/36 (100) | 1 | 24/24 (100) | 12/12 (100) | 12/12 (100) | 1 |

A PCA was then performed to assess the relationship between operon presence and MIC of FDC. More than 90% of the variance in operon presence in the WT/TEM-*Ec* and NDM-*Ec* was respectively explained by the first eight (92.72% of variance) and seven (93.82% of variance) principal components (PC) of the PCA, which were used as factors for the MIC regressions. For the analysis of the WT/TEM-*Ec* collection, the significant factors were PC2 (95%IC = [1.136,2.002], p < 0.001), PC4 (95%IC = [0.316,1.385], p = 0.002), PC5 (95%IC = [0.797,2.072], p < 0.001) and PC6 (95%IC = [0.556,2.052], p = 0.001), which were all positively correlated with the Log_2_ of MIC (Table S2). For the analysis of the NDM-*Ec* collection, the significant factors were PC1 (95%IC = [0.168,0.697], p = 0.002), PC2 (95%IC = [1.097,1.833], p < 0.001), PC5 (95%IC = [0.650,1.972], p < 0.001) and PC7 (95%IC = [0.645,2.542], p = 0.002), which were all positively correlated with the Log_2_ of MIC (Table S3). In both cases, the *fec* operon was the only one whose presence was positively correlated with all the significant components (Table S4 and Table S5). For both collections, the changes in MIC predicted by the model are therefore highly correlated with the presence of the *fec* operon in the isolates (Figure 4).

**Figure 4.**
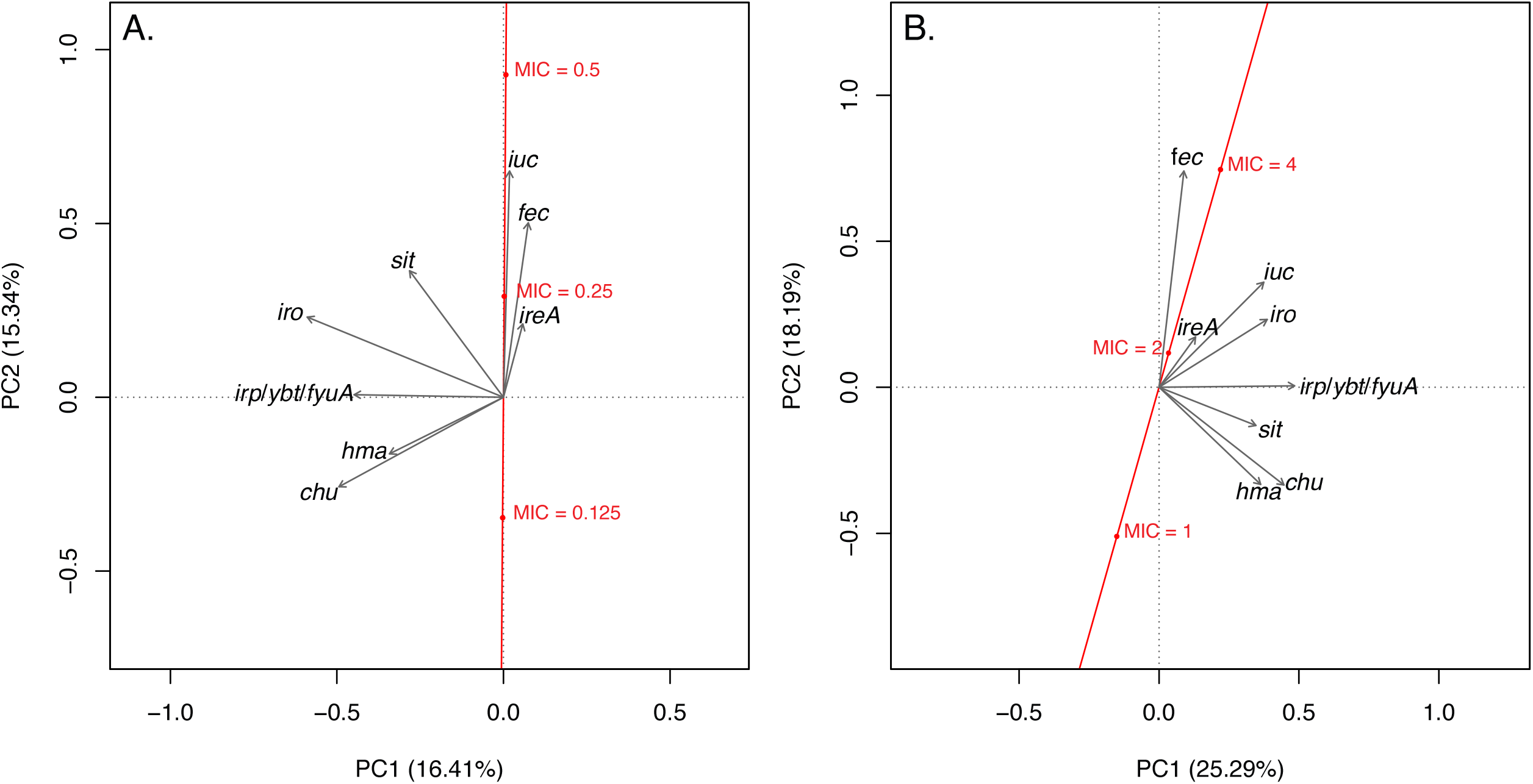
Correlation between the presence of operons and the first two axes (grey arrows) of the principal component analysis of the WT/TEM-*Ec* (A) and the NDM-*Ec* (B) collections, together with the predicted variation of MIC with those two axes (red line), with the 95% confidence interval of the prediction (red envelope). The MIC at the (0,0) coordinate is the average over the whole collection, and the red dots represent the position on the prediction line of measured MIC relative to this average.

### Prevalence and genomic location of *fec* iron uptake system in *E. coli*

By screening the EnteroBase encompassing 70,301 *E. coli* available genomes, 46.8% (32,905/70,301) were found to harbor the *fec* operon (*fecABCDE*). The prevalence of the latter varied by phylogroup, ranging from 3.2% (355/11,173) in phylogroup E to 92.5% (1,481/1,601) in phylogroup C (Figure 5A).

**Figure 5:**
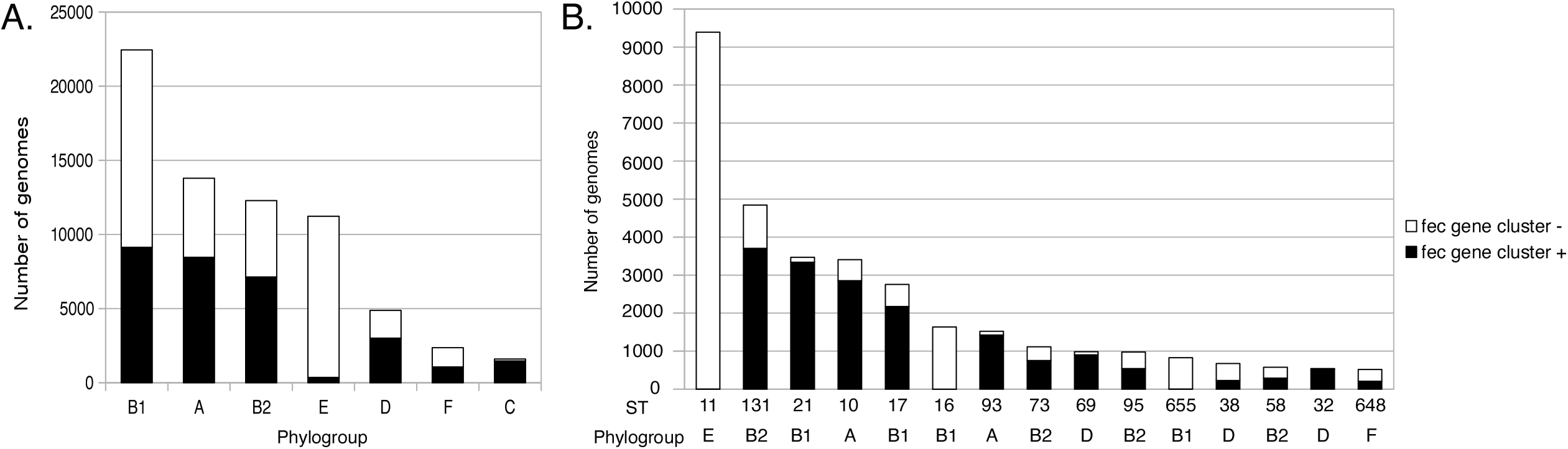
Distribution of the *fec* gene operon among *E. coli* genomes from Enterobase according to their phylogroup (A) and ST (B). Only the 15 most prevalent ST are represented in (B). The number of *fec* gene operon positive and negative genomes are represented by a black bar and a white bar, respectively.

Considering the 15 most prevalent sequence types (STs) in the database, the prevalence of *fec* ranged from 0.1% to 99.4%, with high prevalence observed in both virulent and epidemiologically significant clones from phylogroup B2, such as ST131 (77.0%, 3,729/4,840) and ST95 (55.6%, 545/980), as well as in commensal-associated clones such as ST10 (84.9%, 2,891/3,404) (Figure 5B).

We then focused on a collection of complete genomes from the RefSeq database for the determination of the location of the *fec* operon. Among the 2,302 complete *E. coli* genomes studied, 48.5% (1,116/2,302) were found to harbor the *fec* operon. Most of these operons (99.1%, 1,105/1,116) were located on the chromosome and inserted within a variable genomic region. The complete operon (*fecABCDE*) was present in 99.1% of the chromosom-borne *fec* (1,095/1,106) and 72.7% of the plasmid-borne *fec* (8/11). The majority of incomplete operons contained only the *fecA* gene. Analysis of the eight complete plasmids did not reveal a specific incompatibility (Inc) group associated with the presence of *fec*, although most (4/8) carried the IncFIA replicon.

Additionally, sequences generated by nanopore technology were available for three of the twelve *fec*-positive NDM-*Ec* isolates, enabling analysis of the genomic location of the operon. This *fec* operon was plasmid-located and complete in only a single isolate. In the remaining two strains, the *fec* operon was located on the chromosome.

Further analysis of plasmid-borne *fec* operons suggested distinct origins (Figure S1). The *fecA* gene from our NDM-*Ec* isolate and three plasmids in *E. coli* RefSeq genomes (NZ_CP060942.1, NZ_CP021680.1, NZ_CP026937.1) was closely related to plasmid-borne *fecA* in *Klebsiella pneumoniae* (18). In contrast, *fecA* from other *E. coli* plasmids (n=5) were more closely related to chromosomal *fecA* sequences of *Enterobacter hormaechei*.

## Discussion

In this study, we investigated the role of various *E. coli* iron transport systems on the activity of FDC, using mutants lacking iron capture genes from well-characterized collections of *E. coli* K-12 (Keio knockout collection) and the uropathogenic *E. coli* 536 (536-PAI-deleted collection) as well as genome-sequenced clinical isolates exhibiting different β-lactam phenotypes susceptibility. The latter were either wildtype or producers of TEM penicillinases (WT/TEM-*Ec*) or of NDM-1 carbapenemase (NDM-*Ec*). We confirmed that the two major entry pathways for FDC are the enterobactin receptors CirA and Fiu. The presence of other siderophore such as salmochelin or yersiniabactin had no impact on FDC activity. Interestingly, the presence of the ferric citrate transporter Fec was associated with increased MICs to FDC and contributed to resistance in NDM-producing strains.

*E. coli* possesses a wide array of iron uptake systems, including heme transporters (Chu and Hma), ferrous ion transport systems (Sit, Feo, Efe) associated with non-specific porins (OmpF and OmpC), high-affinity ferric transporters (Fhu, Fec, Kfu) and siderophore systems (enterobactin, salmochelin, aerobactin and yersiniabactin). These siderophore systems, also considered as virulence factors play a central role in iron acquisition under iron-limited conditions (19,20). The catecholate enterobactin siderophore is ubiquitous in *E. coli* and enables Fe^3+^ uptake through its membrane receptors FepA, CirA and Fiu. The activity of FDC is primarily linked to its catechol moiety, which mimics a siderophore and enhances its penetration into the periplasmic space of Gram-negative bacilli (21).

In our study, the FDC MIC of the Δ*cirA* and Δ*fiu* mutants, in iron-deficient media, were significantly higher than that of the K-12-WT, confirming that FDC relies on CirA and Fiu receptors to reach the periplasmic space (22,23). These differences were even more pronounced in iron-supplemented conditions. The stronger impact of *cirA* deletion compared to *fiu* deletion is consistent with previous studies reporting that FDC has much higher affinity for CirA than for Fiu with dissociation constants (Kd) of 63 nM and 0.4 µM respectively (22,24). The lack of effect of *fepA* gene deletion on FDC MICs, under both iron deficient and enriched conditions, also confirmed previous findings indicating the inability of FDC to bind the FepA receptor (22).

This observation aligns with known resistance mechanisms to FDC in Enterobacterales, which often involve mutations in *cirA* (8,10,25–27) or *fiu* (10,25) genes, particularly truncations in *cirA* that can raise MICs higher than 32mg/L. Mutation in the regulatory systems *baeS* and *envZ* which control the expression of *fiu* and *cirA*, have also been associated with increased FDC MICs (27). Thus, the *cirA* and *fiu* gene sequences in each clinical strain were analysed to exclude any mutations affecting FDC susceptibility. Among the resistant isolates, only one strain possessed a truncated *cirA* gene, ruling out bias in our identification of a resistance determinant.

Interestingly, deletion of *fecA*, known as a virulence factor in uropathogenic *E. coli* strains (28), led to a decrease rather than an increase in FDC MIC. This effect was also more pronounced under iron-enriched conditions, with FDC MIC of the Δ*fecA* mutant increased only slightly to 0.08 mg/L whereas MICs of the K-12 WT or other mutants increasing to 0.5-4 mg/L at the highest iron concentration. Furthermore, the *fec* operon was present in all or most clinical NDM-*Ec* and WT/TEM-*Ec* isolates with FDC MICs above or equal to the median MIC, (1.75 mg/L and 0.25 mg/L respectively) but absent or showing low prevalence in isolates with FDC MICs below the median MIC.

The higher prevalence of the aerobactin system (*iuc*) among WT/TEM-*Ec* isolates with FDC MICs above or equal to the median MIC appears to result from a random co-occurrence with the *fec* operon. This association was not observed in the NDM-*Ec* collection, and no difference in MIC was found between *fec*-positive/*iuc*-negative and the *fec*-positive/*iuc*-positive isolates.

Kocer *et al*. previously identified the *fec* operon as a contributor to FDC resistance in clinical isolates producing VIM-type carbapenemases (29). Acquisition of a *fec* operon by a VIM carrying *Citrobacter freundii* led to at least 4-fold increase in FDC MIC, resulting in phenotypic resistance. While the presence of an MBL alone, such as VIM or NDM typically increases FDC MIC by 4 to 6 dilutions compared to the wildtype strains (7,29) which generally remain below the EUCAST clinical breakpoint. However, both the combination of MBL production and *fec* operon could result in a significant FDC MIC increase, exceeding the clinical susceptibility threshold.

Overall, *fec* is widespread in *E. coli*, present in nearly half (46.5%) of the 70,301 strains available in Enterobase. It is found in both epidemiologically significant virulent clones, such as ST131 and ST95, and commensal-associated clones, such as ST10. While commonly located on the chromosome, it can also be found on mobile genetic elements such as plasmids. Plasmid-borne *fec* operons identified in *E. coli* genomes from the RefSeq database were closely related either to chromosomal genes of *Enterobacter hormaechei* or, as observed in one of our NDM-producing isolates, to plasmid-borne genes similar to the recently described plasmid-borne operon from *K. pneumoniae* (18). On one hand, this suggests the mobilization and horizontal from the chromosome through a plasmid vehicle; on the other hand, it indicates the subsequent dissemination of such plasmids across different bacterial species. Consistent with this, in the study by Kocer *et al*. (29), *fec* was carried by a broad host-spectrum plasmid, transferable by conjugation to other *Enterobacterales* but also to *Pseudomonas aeruginosa*. The presence of the *fec* operon on transferable genetic elements poses an additional threat to FDC efficacy on carbapenemase producing strains as FDC remains a last-line therapeutic option.

Several hypotheses may explain the increase of FDC MICs associated with the presence of *fec*. One hypothesis is the downregulation of *cirA* and *fiu* expression by the iron uptake regulator, Fur, in response to increased intracellular iron levels mediated by Fec, as observed by Polani *et al*. (18). However, Kocer *et al*. (10) reported no significant difference in *cirA* and *fiu* transcript level with or without *fec*. Thus, an alternative hypothesis, is that, due to its high affinity for Fe^3+^, the Fec system became the preferred pathway for acquiring iron, thereby reducing the need for siderophores such as enterobactin. This could in turn limit the FDC entry *via* its catecholate moiety, reducing its susceptibility (Figure 6).

**Figure 6.**
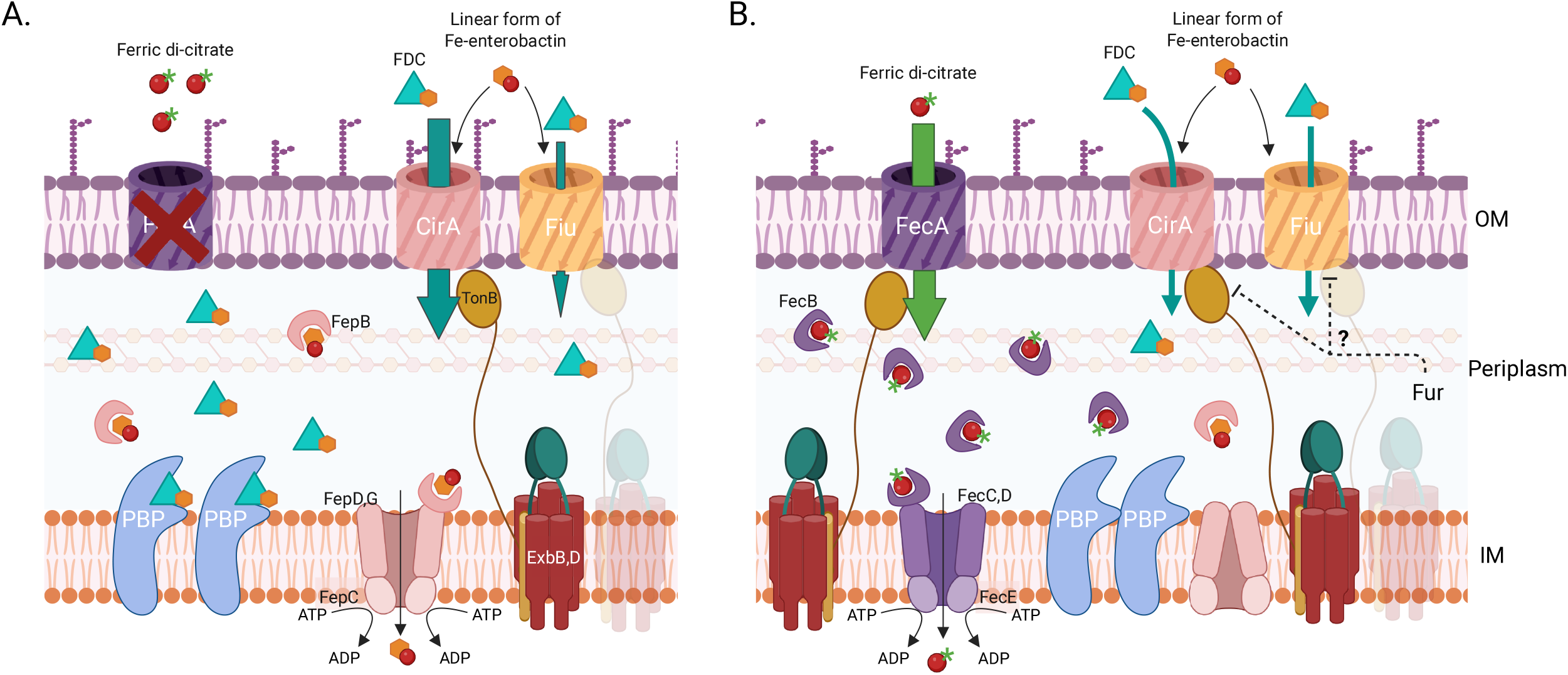
Schematic representation of the hypothesis of the link between operon *fecABCDEIR* prevalence and Cefiderocol (*FDC*) activity. (A) Preferential iron entry *via* the enterobactin (Ent) uptake system in strains not carrying *fecABCDEIR* operon enabling massive FDC entry (low MIC). (B) Preferential entry of iron *via* the FecA transporter in strains which acquired *fecABCDEIR* operon resulting from either the downregulation of *cirA* and *fiu* by Fur or a higher affinity of FecA for Fe^3+^. Thus, this would lead to a reduced use of the enterobactin uptake system and therefore FDC entry (increased/high MIC). Abbreviations: Outer membrane (OM), Inner membrane (IN), Penicillin binding protein (PBP). Created in BioRender. Mallecot, E. (2026) https://BioRender.com/ub5eltm (License GN28XGW79X)

The other siderophore systems, salmochelin and yersiniabactin, investigated using 536-PAI-deleted collection, had no impact on FDC MICs in either iron-depleted or iron-supplemented conditions. This finding suggests that there is no competition between siderophore systems. Finally, no additional associations between FDC susceptibility and the prevalence of iron uptake genes were observed among the collection of clinical isolates.

From an evolutionary perspective, our data provide a new molecular mechanism involving iron, that breaks the classical trade-off between virulence and antibiotic resistance (30) as the strains acquiring *fec* are both more virulent and more resistant. These data are an additional illustration of the many situations in which iron and antibiotics have combinatorial effects when used together, sometimes referred to “liaisons dangereuses” (31).

Our study has several limitations. First, all mutants used (except ΔPAI-III and PAI-dTTP) were single-gene deleted for iron uptake genes, which prevents evaluation of potential additive or synergistic effects between iron uptake systems, particularly in combination such as *cirA/fiu, fec*/*cirA* or *fec*/*fiu* double mutants. Second, we determined only FDC MIC and did not performed time-killing assays, as conducted in other studies (32). Finally, we did not investigate gene expression or protein levels. Therefore, we are unable to state whether *fec* has an impact on the regulation of *cirA* and *fiu* expression, in view of the conflicting results published by Polani *et al*. (18) and Kocer *et al*. (29). Thus, the mechanism behind FDC MICs rise in presence of *fec* need to be investigated in further analysis.

## Conclusion

In addition to confirming the role of the CirA and Fiu receptors, our study highlights the ferric citrate transporter Fec as a novel resistance determinant affecting FDC activity in *E. coli*. This is particularly concerning given its prevalence in nearly 50% of *E. coli strains*, its potential for plasmid-mediated transfer, and its ability to confer FDC resistance when combined with other mechanisms, such as MBL production. By linking increased virulence and resistance, the *fec* operon exemplifies how iron acquisition can override the classical trade-off between pathogenicity and antibiotic susceptibility.

## Acknowledgment

We thank Professor Katell Peoch of the biochemist laboratory of Bichat-Claude Bernard (BCB) University Hospital for providing the ammonium iron (III) citrate used in our study.

